# Non-conscious Multisensory Integration in the Ventriloquist Effect

**DOI:** 10.1101/2025.06.01.657322

**Authors:** Elise Turkovich, Soorya Sankaran, Wei Dou, Jason Samaha

## Abstract

The degree to which information from distinct sensory modalities can interact in the absence of conscious awareness remains controversial. According to the Global Neuronal Workspace Theory (GNWT), unconscious sensory information remains relatively confined to sensory cortex and should not be capable of interacting with other modalities until it is broadcast into the (conscious) global workspace comprising late (>300ms) frontal-parietal activation. The ventriloquist effect is a classical multisensory integration phenomenon that refers to the misperception of sound location towards concurrent visual stimulation, such as perceiving the voice of a ventriloquist actor as coming from the moving dummy. Here we used meta-contrast masking to render a brief flash stimulus non-conscious while 30 participants (male, female, and non-binary) performed a sound localization task. We found that, despite being subjectively invisible and at chance at discriminating the flash location, participants were nevertheless biased to localize sounds towards the unconscious flash locations. The effect was present in virtually all participants, was nearly as large as the effect on conscious trials, and was robust to controls for individual differences in task performance. Decoding and ERP analyses of concurrently recorded EEG signals showed that the non-conscious flash location information was present up until around 220ms but not after; suggesting that the visual influence on sound perception may have occurred before conscious broadcast. In support of this, early decoding accuracy on non-conscious trials predicted individual variation in the size of the unconscious ventriloquist effect. Our findings suggest that subjective perception is not required for the integration of signals originating in distinct sensory modalities prompting new questions about the role of subjective perception in multisensory integration.

## Introduction

The role that awareness plays in cognitive and perceptual functions is a central topic in the development of theories of consciousness (Block, 1995; Cohen & Dennett, 2011; Ludwig, 2023; Samaha, 2015; Seth & Bayne, 2022). Most current neuroscientific theories of consciousness propose that conscious perception confers some functional advantage that non-conscious processing lacks (Lau & Rosenthal, 2011; Seth & Bayne, 2022). One proposed function, common to many approaches, is that consciousness has a role in integrating neural processes that would otherwise be independent (the so-called “integration consensus” (Morsella et al., 2016). A prime example of the proposed integrative function of consciousness is in the global (neuronal) workspace theory (GNWT) of consciousness (Baars, 2005; Dehaene & Changeux, 2011; Dehaene & Naccache, 2001; Mashour et al., 2020), which posits that non-conscious processing remains relatively siloed within specialized processing modules where it is inaccessible to other cognitive, mnemonic, motor, evaluative, and perceptual processes.

Consciousness, on the other hand, reflects the contents that are broadcast beyond local processing and are widely accessible by other processing modules, putatively mediated by a long-range, recurrent, frontal-parietal network (Mashour et al., 2020). Such broadcast is thought to happen relatively late (around 250-300 ms post-stimulus) in a non-linear, “all-or-none” fashion, and to underlie a subject’s ability to report their perceptual experiences (Dehaene et al., 2014; Del Cul et al., 2007; Salti et al., 2015; van Vugt et al., 2018).

One prediction that follows from the integration function proposed by GNWT theory is that unconscious processing in distinct sensory modules should not interact. Or, as Baars puts it *‘consciousness is needed to integrate multiple sensory inputs’* (Baars, 2002 p. 47–48). Several lines of evidence support the idea that non-conscious sensory processing remains local and confined to the cortical areas selective for the type of sensory information. For example, visual stimuli that are reported as unseen by the observer elicit brain responses in early and mid-level visual areas but typically not in non-visual areas or higher-order association areas (Dehaene et al., 2001; Del Cul et al., 2007; Sergent et al., 2005). Non-conscious processing of auditory stimuli does not elicit spiking activity in somatosensory areas (Noel et al., 2019) and does not lead to global broadcast (Sadaghiani et al., 2009) particularly, when also task-irrelevant (Sergent et al., 2021). In contrast to this prediction, however, a growing number of studies point to conditions under which integration can occur non-consciously (Mudrik et al., 2014; but see Biderman & Mudrik, 2018; Hirschhorn et al., 2020) even when both of the to-be-integrated stimuli are non-conscious and come from different sensory modalities (Scott et al., 2018).

However, research examining audio-visual speech processing in relation to conscious perception has generally found that, while congruency between one conscious modality (e.g., speech sounds) and another unconscious one (e.g., movies of faces rendered invisible through inter-ocular suppression techniques) can produce priming effects on subsequent response times (RT) (Ching et al., 2019; Palmer & Ramsey, 2012; Plass et al., 2014) or on suppression breakthrough times (Alsius & Munhall, 2013) genuine cross-modal integration as reflected in the illusory percept of a McGurk effect has, so far, not been demonstrated unconsciously (Ching et al., 2019; Palmer & Ramsey, 2012).

A canonical example of multisensory integration is the Ventriloquist effect (VE), whereby the location of a visual stimulus biases the perceived location of a concurrent auditory stimulus, such as when perceiving the moving mouth of a ventriloquist dummy as the spatial source of vocalization (Alais & Burr, 2004; Vroomen & De Gelder, 2004). Recent evidence suggests that visual stimuli which are not consciously perceived may nevertheless bias the perceived location of auditory stimuli. Delong et al., (2018) used dynamic continuous flash suppression (dCSF) to render a brief flash invisible while having subjects judge the spatial location of a concurrent white noise burst. They found a reliable VE across two experiments, suggesting that the non-conscious flash still biased the perceived sound location (Delong et al., 2018). However, one potential drawback of the use of dCSF for masking in this context is that subjects may have actually perceived the brief flash stimulus consciously (or part of it) but simply confused it for a part of the mask, given the visually complex and dynamic appearance of the Mondrian patterns used, which varied over space and time in color, motion, and luminance, among other features (Gelbard-Sagiv et al., 2016). Additionally, prior studies have not yet investigated the timecourse of neural processing underlying putative non-conscious integration in the VE.

We conducted a multi-day behavioral and EEG experiment using meta-contrast masking to render brief flashes invisible while subjects judged the location of a concurrent auditory stimuli. Using both objective criteria for awareness (chance performance in flash localization) and subjective criteria (reports of “no experience” on the perceptual awareness scale; PAS), we found a strong non-conscious VE that was present in virtually every subject. Decoding location information from concurrently recorded EEG signals revealed that non-conscious representations of the visual stimulus location were present only for an initial 220 ms, consistent with a lack of late global broadcast, suggesting that non-conscious audiovisual integration may occur prior to 220 ms.

## Methods

### Participants

32 members of the UC Santa Cruz community participated in this two-day study (age range 18-32, 18 Female, 12 Male, 2 Non-Binary). All subjects were compensated with a $20 Amazon gift card and class credit when appropriate. One subject was excluded from analyses due to excessive noise in the EEG signal and another due to blinking in response to most auditory stimuli, leaving 30 subjects in the final analyses (unless otherwise noted). The study was approved by the UC Santa Cruz Institutional Review Board.

### Stimuli

The experiment was implemented in MATLAB (The MathWorks Inc., 2022) and all stimuli were generated using the Psychophysics Toolbox Version 3 (Kleiner et al., 2007; Pelli, 1997). Stimuli were presented on a uniform gray background (approx. 50 cd/m^2^) on a gamma-corrected VIEWPixx EEG monitor (53.4 × 30 cm, 1920 × 1080 resolution, 120 Hz refresh rate). Participants were seated in a dimly lit room approximately 74 cm away from the screen with their head stabilized on a chin rest. Auditory stimuli were delivered using Etymotic Research 3C insert earphones at an (individually adjusted) comfortable volume.

To deliver auditory stimuli originating from varying spatial locations, the MIT KEMAR head related transfer function (HRTF) was convolved with the auditory stimulus (Gardner & Martin, 1995). Specifically, the timing and amplitude profile of a 20ms burst of white noise (generated randomly on each trial) was manipulated for each ear to simulate sound from 5 deg. to the left, center, or 5 deg. right, along the azimuth with zero elevation.

The target visual stimulus was a circular luminance increment subtending 1 deg. of visual angle, whose luminance contrast with respect to the gray background was individually determined in an initial thresholding procedure (see below). The masking stimulus was an annulus with an outer diameter of 2 deg., an inner diameter of 1 deg., and a luminance of ~98 cd/m^2^. A central crosshair plus bullseye (0.6 deg.) was used as a fixation point

### General Procedure

We modeled our task design closely on that of Delong et al., (2018) with the primary difference being that we used meta-contrast masking rather than dCFS to render visual stimuli invisible. Participants completed this experiment over a two-day period (see Figure 1). On the first day, lasting approximately one hour, participants practiced the auditory localization task (described next) until they achieved above-chance performance. Then participants completed the visual thresholding task, designed to ensure at or close to chance performance on the visual localization task. Both of these tasks were in preparation for the main task on Day 2, during which we aimed for participants to be proficient at auditory localization but unaware of the visual stimulus on a majority of trials. On Day 2 (which occurred within one week of Day 1), participants were fitted with EEG, completed one more practice block of the auditory localization task, a visual only block, eight blocks of the main AV task, and then a final visual only block. Each of these tasks is described in greater detail below. Only data from Day 2 is analyzed here.

**Figure 1.**
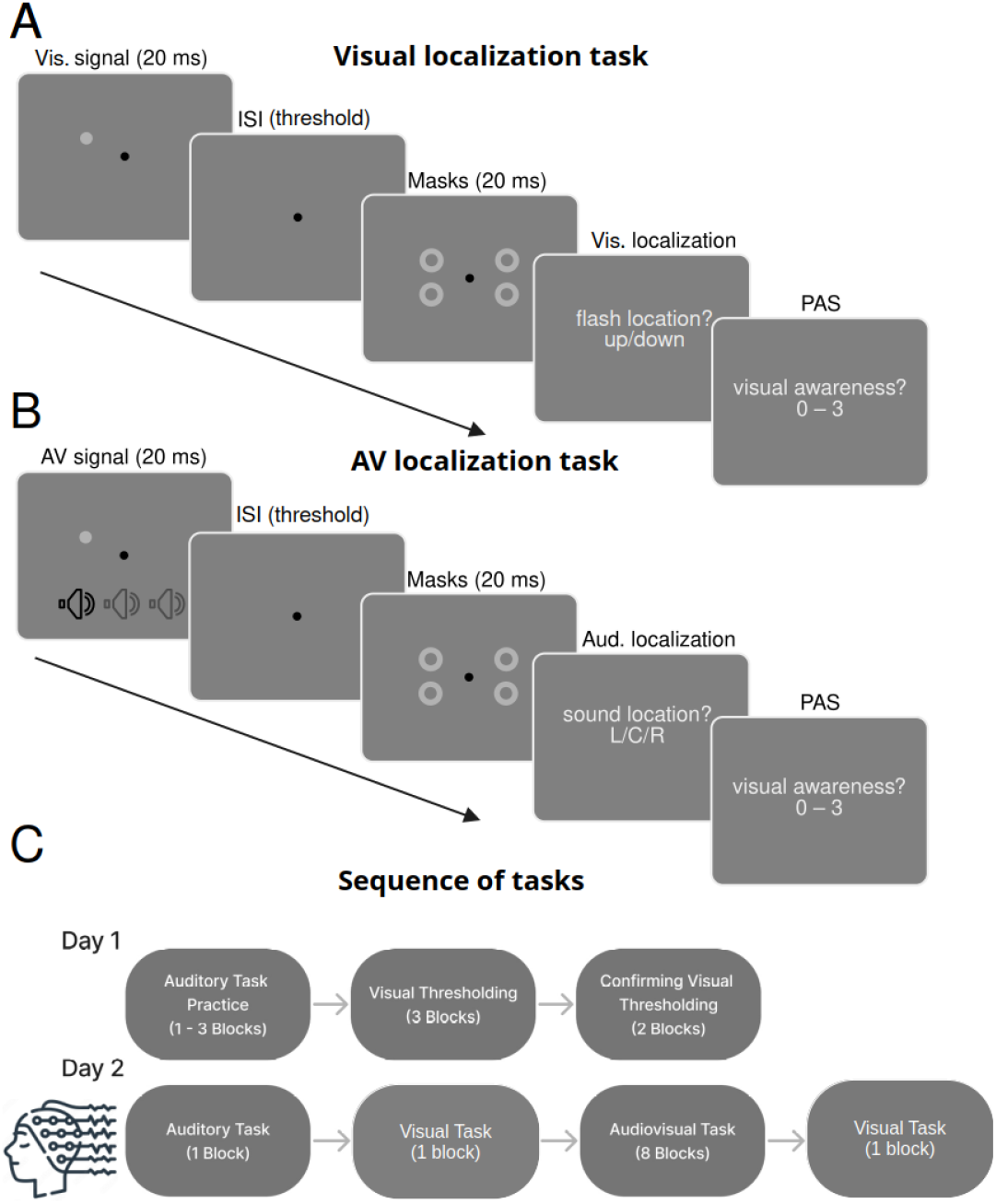
Tasks and procedure. **A)** In the unimodal visual localization task, participants judged the vertical location (up or down) of a brief flash target and then rated subjective visibility on the 4-point Perceptual Awareness Scale (PAS). Targets were backwards-masked with metacontrast annuli that appeared at all possible target locations in each visual quadrant. The target luminance and inter-stimulus interval (ISI) between mask and target were determined in a separate thresholding session so as to produce near chance localization accuracy on trials judged as invisible (PAS = 0). **B)** In the main audio-visual (AV) localization task, participants were shown threshold flash stimuli concurrently with supra-threshold auditory stimuli presented at three simulated locations across the horizontal plane (using HRTFs; see methods)> Participants first judged the auditory stimulus location (left, center, or right) and then flash visibility using the PAS. Visual and auditory locations were randomized so that one did not inform the location of the other. **C)** Sequence of tasks across two days of testing. On Day 1 (behavior only) participants practiced auditory localization (in the absence of visual stimuli), then performed several blocks of visual thresholding in order to find the ISI and flash luminance that achieved at or near chance performance on trials rated as subjectively invisible. On Day 2 EEG was recorded while subjects performed a pre-task visual-only block to check for chance performance on invisible trials, the main audiovisual task, and then a post-task visual-only block.

### Auditory localization task

In the auditory localization task, the participant was presented with a 20ms burst of white noise originating from 5 deg. left, center, or 5 deg. right. The task was to judge the sound’s location using one of three options, ‘left’, ‘center’, or ‘right’, using the keys 7, 8, and 9, respectively, with the right hand. An inter-trial-interval (ITI) randomly sampled between was 0.9 and 1.4 sec followed the response. Given the difficulty of discriminating sound locations separated by only 5 deg., we repeated blocks of 72 trials with visual feedback displaying the correct answer on every trial until each participant performed higher than 45% accuracy across an entire block (where chance is 33%). This typically took between 1-3 blocks, all performed on Day 1. A single block was also run at the beginning of Day 2 to remind participants of the auditory judgment before proceeding to the rest of the experiment.

### Visual thresholding task

The visual thresholding task was designed to find the ideal masking conditions for rendering the flash invisible on a majority of trials. To establish the optimal inter-stimulus interval (ISI) between the target flash and the mask for each participant, 3 blocks (576 trials total) of a purely visual masking task were conducted on Day 1. The visual task involved fixating while a brief (20ms) flash was presented at one of four quadrant locations, 1.2 deg. above or below and 5 deg. left or right from fixation. After a brief ISI, a mask was presented for 20ms in all four quadrants, encompassing the space around all potential flash locations. The ISI varied between 0-40ms (corresponding to SOAs of 20-60ms), in steps of 8ms. While doing the task, participants are asked 1) to rate their perceptual awareness of the target stimulus on the 4-point perceptual awareness scale (PAS; Sandberg et al., 2010) and 2) to discriminate the vertical location of the stimulus (i.e., upper or lower visual field). The numbers 0-3 of the PAS correspond to ‘not seen’, ‘brief glimpse’, ‘almost clear’, and ‘clear image’, respectively.

Participants were repeatedly instructed to use a PAS response > 0 if they had any feeling whatsoever of having seen the stimulus. The PAS response aims to capture the participants’ subjective invisibility on a trial-by-trial basis, i.e., does the participant believe they saw the flash? The location discrimination question aims to provide an objective measure of stimulus invisibility based on the reasoning that when location discrimination accuracy is indistinguishable from 50% (i.e., chance), the participant is likely unaware of the visual stimulus (Merikle et al., 2001).

At the end of all three thresholding blocks on Day 1, plots were produced displaying accuracy at discriminating the flash location at each ISI. The timing at which the participant was closest to chance on trials rates as invisible is selected. If no ISI led to an accuracy level below 60% in invisible trials, the luminance of the flash was reduced from a starting luminance of ~70 cd/m^2^ to ~60 cd/m^2^ and the procedure was run again. Two subsequent shorter blocks (96 trials) with the fixed ISI and luminance from the thresholding were then completed to confirm the decision and modify parameters slightly if needed (i.e., make small reductions in luminance if performance was greater than 60%), all performed on Day 1

### Visual Localization Task

The masking parameters from the Day 1 thresholding procedure were fixed for all subsequent visual tasks in Day 2. Specifically, on Day 2, participants performed two blocks (48 trials each, one before and one after the main AV task) of a visual only task identical to the visual thresholding task but with a constant ISI and flash luminance. The purpose was to test that performance was still near chance on trials rated as invisible.

### Main Audio-Visual Task

The main task involved the presentation of a temporally congruent audio-visual stimulus with varying spatial locations. On Day 2, while collecting EEG data, participants complete eight blocks of the combined AV task (720 trials total), wherein the flash target was presented in temporal congruence with the sound burst, followed by the visual masks. Participants are asked (1) The location of the sound (left, center, or right) and (2) their perception of the visual stimulus on the PAS scale (0-3). The visual location judgment was omitted from this main task based on pilot data where subjects indicated occasionally confusing response options and difficulty remembering all responses when asked three questions per trial.

### EEG Recording and Pre-Processing

EEG data were recorded at 1000 Hz using a 64-channel Ag/AgCl gel-based active electrode system (actiCHamp Plus, Brain Products) with electrode FCz as the online reference. The data were pre-processed using custom MATLAB scripts (version R2023a) and the EEGLAB toolbox (Delorme & Makeig, 2004). First, a band-pass filter was applied at from 0.1 – 50 Hz (zero-phase, Hamming-windowed sinc FIR filter), the data were downsampled to 500 Hz, and then re-referenced to the median of all electrodes. Epoched data were then visually inspected and trials with excessive noise, muscle artifacts, or ocular artifacts overlapping with the stimulus presentation were rejected. Independent components analysis (ICA) was performed using the INFOMAX algorithm to remove 1 or 2 components per subject corresponding to blinks or eye movements. An average of 55 trials were removed (range: 10-167). Channels with excessive noise were removed and spherically interpolated (average of 1.4 channels; range: 0-6). Lastly, the data were re-referenced to the average across channels and a 200 ms pre-stimulus baseline was subtracted from each timepoint, channel, and trial.

### Behavioral Analysis

All behavioral analyses were conducted on data from day two, which comprised the auditory-only task, the main AV task, as well as the two visual-only blocks conducted immediately before and after the main AV task (see Figure 1). We categorized trials as invisible when PAS=0 and visible when PAS>0. We computed a ventriloquist effect (VE) index by considering the mean sound localization report in deg. (i.e., −5, 0, 5) as a function of the actual sound location (in deg.) and the flash location (left or right or absent). We computed the VE (in deg.) as the difference in mean sound localization report between trials with a left or right flash. Accordingly, if the flash location had no influence on sound localization, the VE would be 0 and if the flash attracts (repels) the sound localization report, the VE is positive (negative). Thus, the classic VE would correspond to a positive value using this metric. We tested whether an unconscious VE occurred by computing the VE index separately for invisible (PAS=0) and visible (PAS>0) flash trials and comparing each index to 0 using repeated-measures t-tests. Bayes factors were also used to evaluate the evidence for the null (BF_null_) or alternative (BF_alt_) hypothesis using the Bayes Factor MATLAB package from Bart Krekelberg (https://github.com/klabhub/bayesFactor), which uses a JZS prior.

### EEG Decoding Analysis

We used multivariate pattern analysis of all EEG channels to decode stimulus information from brain responses in order to study the timecourse of neural representations of stimulus location. We first sought to decode visual location information in order to track how neural processing of the flash changed with the subject’s awareness. Based on the predictions of GNWT theory, we hypothesized that the flash location on invisible trials may be decodable early on (reflecting putatively unconscious processing) but should not be “broadcast” to the fronto-parietal workspace supporting sustained representation and thus should no longer be decodable after the time broadcast is assumed to occur (around 250 to 300 ms). To this end, we trained a linear discriminant analysis (LDA) classifier to predict the horizontal location (left or right) of the flash stimulus during the main AV task. We focused on the horizontal dimension of the visual stimulus since it is the relevant dimension along which the perceived auditory location might be biased. Single-trial EEG data from all sensors were first downsampled to 250 Hz to speed up computation, z-scored across time, and under-sampled to ensure an equal number of left, right, up, and down flash trials at each of the three sound location so as not to bias classification by any imbalance in either the flash or sound locations. Classification was performed using 5-fold cross validation repeated a total of 30 times per participant (each time using a different random under-sampling of equated trial-types so as not throw data out). We then averaged classification accuracy by visibility rating (invisible, PAS = 0; visible PAS > 0) and then across the 30 repetitions for each participants. Because some subjects had too few trials either rated as visible or invisible for stratification into 5 folds, we restricted all decoding analysis to subjects with at least 30 trials in the visible and 30 trials in the invisible conditions. Additionally, to ensure that decoding on invisible trials was not driven by above-chance objective performance in some participant’s behavior, we further excluded subjects from decoding if Bayesian analysis did not indicate evidence in favor of the null hypothesis of chance performance on invisible trials (i.e., if the BF_null_ of a binomial test of invisible accuracy against 0.5 was below that 3.33; see *Behavioral Control Analysis* section in the *Results*). Together, these constraints left 20 subjects for decoding. After sub-sampling, the mean (SEM) number of trials included in the visible and invisible decoding was 173 (20.5) and 175 (18.4) respectively. Decoding was implemented in MATLAB using the MVPA-light toolbox (Treder, 2020).

To statistically evaluate decoder performance, we tested whether classification proportion correct at each time point differed from chance levels (0.5) and whether classification differed between visible and invisible trials using two-tailed repeated-measures t-tests. Cluster correction using the cluster size method (Maris & Oostenveld, 2007) was used to correct for multiple comparisons across time. Specifically, the mapping between data and condition was randomized 5,000 times and all timepoints were tested for each permutation. The size of the largest cluster of contiguous significant timepoints was stored for each of the 5,000 permutations and only cluster sizes in the real data that exceeded the 95 percentile of this distribution of cluster sizes expected under the null hypothesis were considered statistically significant.

To visualize the scalp channels contributing to decoding we also ran a searchlight decoding procedure whereby the same decoding pipeline was applied in a moving spatial window centered on each electrode and extending to include about 6 additional neighboring electrodes (fewer at the edges of the cap). The decoding score using just the electrodes in that spatial window was then assigned to the central electrode. We performed this searchlight procedure using data from all timepoints in an “early” window from 170 to 200 ms and a “late” window 230 to 360 ms, based on the temporal profile of the decoding from all electrodes. Separate decoders were run for visible and invisible trials.

We also attempted to decode the sound location (left/center/right) using the same approach described above, however the decoder never reached significant above-chance classification at any timepoint when training and testing on all AV trials together so we did not further explore sound location decoding.

### Lateralized ERP analysis

Lastly, we analyzed lateralized ERPs in response to the different visual stimulus locations as another way to track stimulus-specific processing on visible and invisible trials. To this end, we computed ERPs from electrodes contralateral to the visual stimulus location (averaging channels PO8, P6, and P8 when the flash was on the left and channels PO7, P5, and P7 when the flash was on the right) as well as from electrodes ipsilateral to the stimulus (PO8, P6, and P8 on right flash trials and PO7, P5, and P7 on left flash trials). We then computed the contra-ipsi difference waveform to isolate spatially-specific responses as well as subtract out any common mask-related activity. Difference waveforms were computed separately for visible and invisible trials and compared against each other using the same cluster-correction procedure described above in the decoding analysis. The same 20 subjects used in the decoding analysis were used in the ERP analysis. As in the decoding analysis, we also subsampled trials so that difference waveforms were always based on an equal number of left/right/up/down flash locations and left/center/right sound location trials. This random under-sampling was repeated 30 times per participant and visibility level and averaged over all 30 repetitions. This ensured that the differences between visible and invisible trials were not driven by any possible visual or auditory stimulus differences.

## Results

### Unimodal Visual Localization and Awareness

We first determined whether flash localization accuracy was at chance levels on trials rated as invisible (i.e., PAS = 0; see Figure 2). In the pre-task visual-only block, we found that mean (*SEM)* flash localization accuracy on invisible trials was 50.6% (1.75%) and did not differ from chance (*t*(29) = 0.377, *p* = 0.708) with a BF indicating substantial evidence for the null (BF_null_ = 4.816). Likewise, in the post-task visual only block, mean accuracy on invisible trials was 53.7% (2.27%), which did not differ from chance performance (t(29) = 1.650, *p* = 0.109), with a BF providing only inconclusive evidence in favor of the null (BF_null_ = 1.534). Comparing the two, invisible flash localization accuracy measured pre- and post-task did not differ (*t*(29) = −0.961, *p* = 0.344, BF_null_ = 3.371), suggesting that the thresholding procedure produced stable performance over the course of the experiment. Flash localization accuracy on trials judged as visible, however, was significantly above chance in both the pre-task block (M = 60.7%, SEM = 3.47%, *t*(29) = 3.100, *p* = 0.004, *BF*_*alt*_ = 9.308) and in the post-task block (M = 64.3%, SEM = 3.32%, *t*(29) = 4.318, *p* = 0.004, *BF*_*alt*_ = 163.041), with no difference between the two (*t*(29) = −0.634, *p* = 0.531, BF_null_ = 4.272). Together, this indicates that performance was stable across the experiment, was better than chance when subjectively aware, and was not better than chance when subjectively unaware.

**Figure 2.**
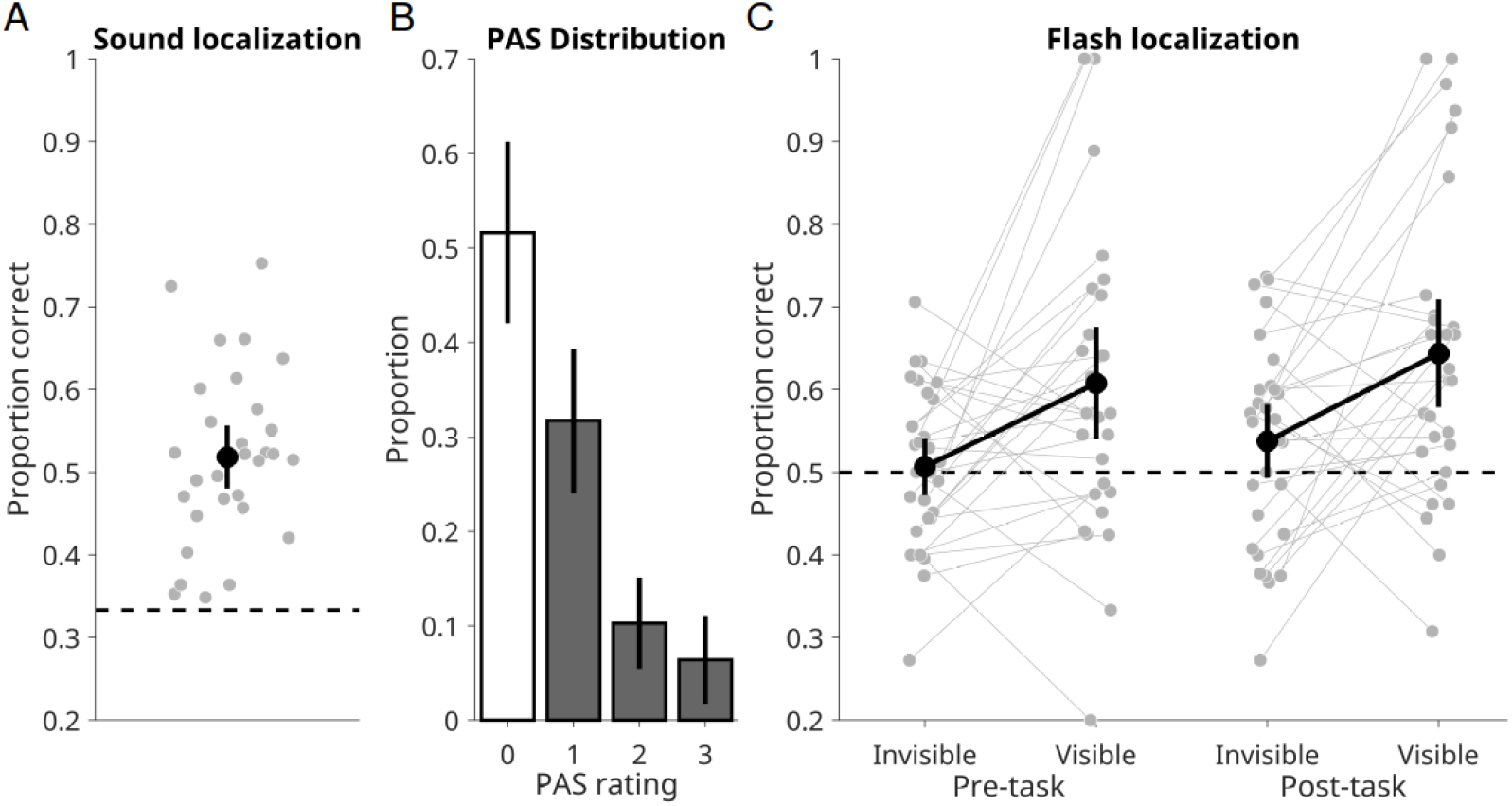
Unimodal visual and auditory behavior on Day 2 (n = 30). **A)** Sound localization accuracy during the main task. All subjects could localize the sounds better than chance levels (0.33). **B)** The distribution of perceptual awareness scale (PAS) ratings in response to the flashes in the main task. All trials rated as 0 on the PAS (“no experience”, white bar, ~ 50% of trials) were considered invisible for the remaining analysis. Any rating above 0 was considered visible. The second most frequent subjective response was 1 or “brief glimpse” **C)** Unimodal visual localization accuracy on trials judged as subjectively invisible (PAS = 0) and visible (PAS > 0) measured immediately pre- and post-main task. Error bars show 95% confidence intervals which overlap chance performance (0.5) on invisible trials, indicating that subjects could not, on average, localize the visual stimuli better than chance levels when they were subjectively invisible. Visible trials, on the other hand, were localized significantly better than chance. All error bars are 95% confidence intervals.

### Unimodal Auditory Localization

We next checked that auditory localization was above chance performance (Figure 2), which ensures that participants were using the timing and amplitude cues in the stimuli to localize the sounds. Indeed, average auditory localization accuracy during the main task was 51.8% (1.91%), well above the chance level of 33% (*t*(29) = 9.664, *p* < 0.001, BF_alt_ > 999).

### Conscious and Non-Conscious Ventriloquist Effect

As seen in Figure 3A, when considering just the visible trials, the presence of the flash on the left or right side of fixation caused sound location judgments to be shifted to the left and right, respectively, relative to flash absent trials (for descriptive statistics of the full distribution of auditory responses across all trial types see Table 1).

**Table 1.** Auditory localization and visibility response percentages for each stimulus type during the main AV task (n = 30)

| Stimulus condition | % left response | SEM | % center response | SEM | % right response | SEM | % visible | SEM |
| --- | --- | --- | --- | --- | --- | --- | --- | --- |
| A<sub>-5</sub> | 46.42 | 2.78 | 34.12 | 2.71 | 19.45 | 2.26 | 33.04 | 5.14 |
| A<sub>0</sub> | 20.66 | 2.2 | 55.37 | 3.57 | 23.97 | 2.54 | 35.26 | 5.07 |
| A<sub>+5</sub> | 13.16 | 1.47 | 30.6 | 2.46 | 56.24 | 2.93 | 35.96 | 5.02 |
| A<sub>-5</sub>V<sub>-5</sub> | 59.3 | 2.82 | 26.6 | 2.53 | 14.1 | 1.69 | 44.34 | 5.03 |
| A<sub>0</sub>V<sub>-5</sub> | 35.54 | 2.99 | 49.16 | 3.46 | 15.29 | 2.13 | 45.55 | 4.98 |
| A<sub>+5</sub>V<sub>-5</sub> | 22.04 | 2.96 | 29.32 | 2.59 | 48.63 | 2.95 | 46.84 | 5.19 |
| A<sub>-5</sub>V<sub>+5</sub> | 35.87 | 2.73 | 29.42 | 2.42 | 34.71 | 2.87 | 48.18 | 5.12 |
| A<sub>0</sub>V<sub>+5</sub> | 12.89 | 1.78 | 48.25 | 3.46 | 38.86 | 3.32 | 51.41 | 5.01 |
| A<sub>+5</sub>V<sub>+5</sub> | 7.69 | 1.26 | 23.82 | 2.29 | 68.49 | 2.7 | 54.01 | 5.08 |

**Figure 3.**
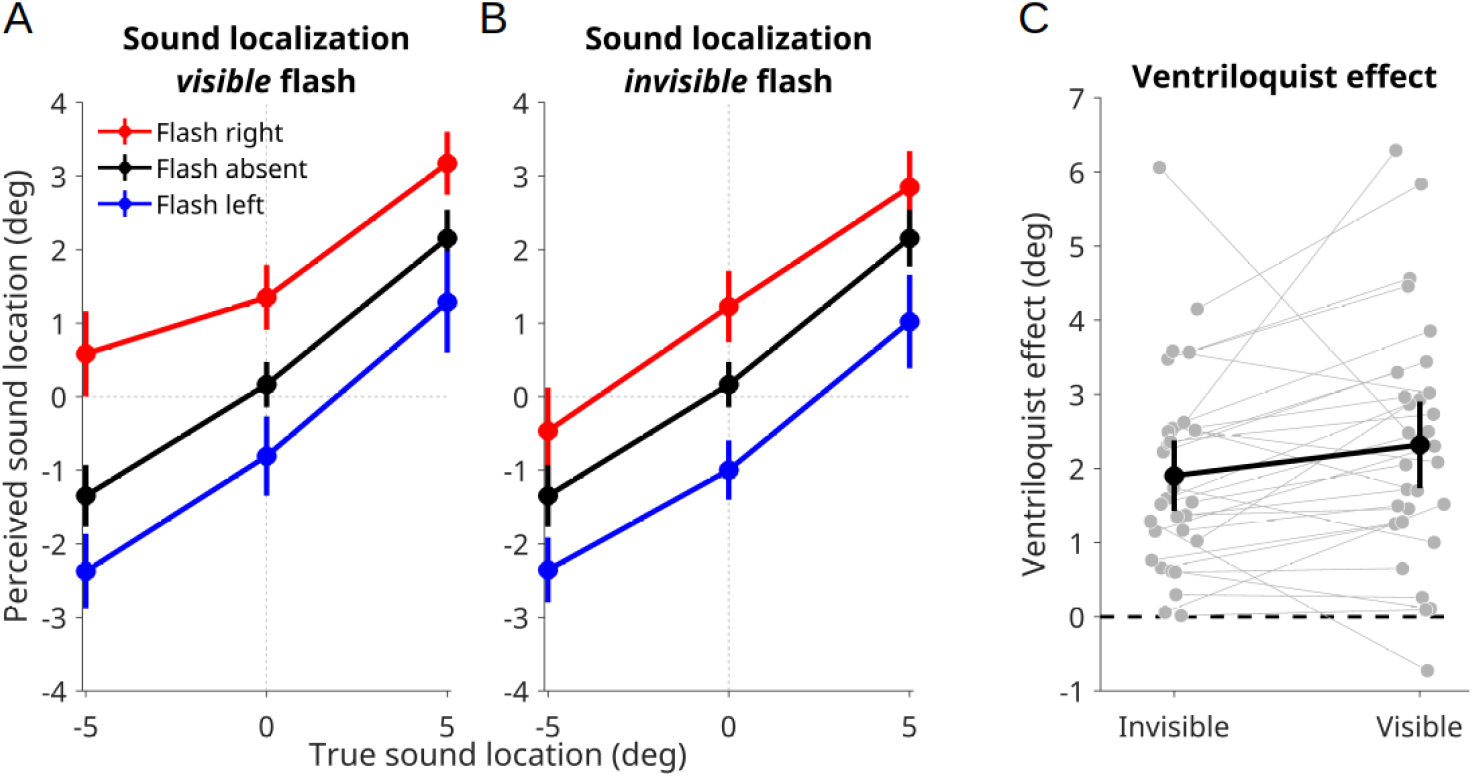
Ventriloquist effect (VE) for visible and invisible trials (n = 30). **A)** Mean reported sound location (left = −5, center = 0, right = +5) as a function of the actual (simulated) sound location for flash absent (black), flash right (red), and flash left (blue) trials when the flash was judged as visible (PAS > 0). Reports generally followed the true sound location but the rightward and leftward shift of sound localization on right and left flash trials respectively indicates a VE, which was quantified as the difference in mean sound localization report on right and left flash trials. Error bars are 95% confidence intervals. **B)** Same as panel A but for trials where the flash was judged as invisible (PAS = 0). Note that the “flash absent” data are the same in both panels A and B and are plotted largely for reference. **C)** The VE index separately for trials where the flash was rated invisible and visible. Error bars are 95% confidence intervals and do not overlap zero in either visible or invisible conditions, indicating a significant VE for subjectively unseen stimuli.

Strikingly, a very similar pattern holds on invisible trials (Figure 3B), with the invisible flash seemingly impacting sound localization. The difference in mean auditory localization reports between left and right flash trials provided an index of the VE, which we further broke down by trials rated as visible (PAS > 0) and invisible (PAS = 0; Figure 3C). On visible trials, we observed a clear VE with visual stimuli biasing auditory localization towards them by an average of 2.31 deg. (0.29), which was significantly different than zero (*t*(29) = 7.776, *p* < 0.001, BF_alt_ > 999). Regarding our main hypothesis, we observed a VE of 1.90 deg. (0.24) even on invisible trials, which represents decisive evidence against the null hypothesis (*t*(29) = 7.799, *p* < 0.001, BF_alt_ > 999).

Although the conscious VE was numerically larger than the non-conscious effect, the two were not significantly different (*t*(29) = −1.747, *p* = 0.091), however, the strength of evidence for the null was inconclusive (BF_null_ = 1.335). Together these results suggest that subjective awareness of a visual stimulus is not needed for multisensory integration and, under conditions of meta-contrast masking, the unconscious VE can be nearly as large as the conscious effect.

### Behavioral Control Analyses

While at the group level participants were not better than chance at localizing the visual stimulus on invisible trials, it is always possible that certain individual subjects were significantly above chance and are driving the non-conscious VE. We addressed this in two ways. First, if this were the case one may expect a positive correlation between an individuals localization accuracy on invisible trials (here, collapsing across pre- and post-task blocks) and the size of their VE on invisible trials. However, a regression analysis revealed no such effect (slope = 0.018, SEM = 0.038, *p* = 0.644; see Figure 4) which a Bayesian correlation indicates is substantial evidence for the null (BF_null_ = 6.36). Moreover, as seen in Figure 4, the 95% confidence interval on the regression line at chance performance on the x-axis does not include a VE of zero, indicating that even for objective flash unconsciousness (p(correct) = 0.5), the predicted size of the VE is around 1.88 deg.

**Figure 4.**
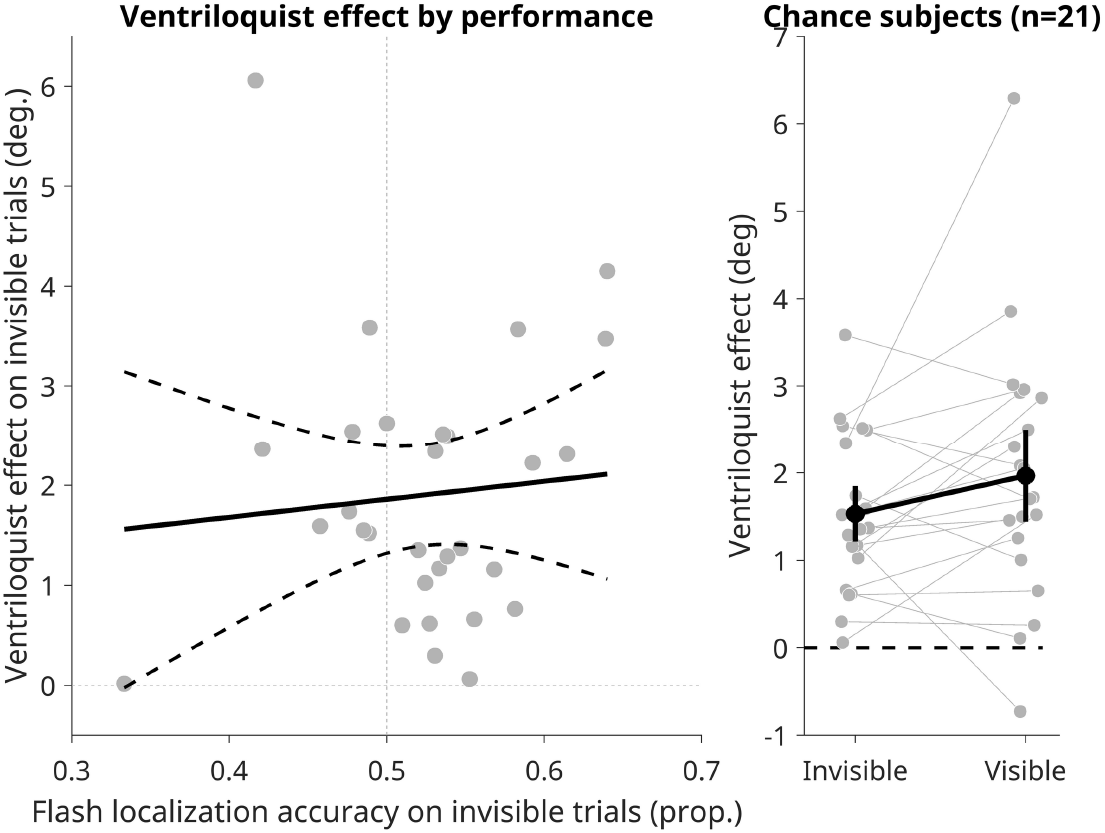
Non-conscious VE does not depend on task performance. The left panel shows each participant’s VE on invisible trials as a function of their flash localization accuracy on invisible trials. A lack of clear correlation and a significant intercept term at p(correct) = 0.5 on the x-axis suggests that the non-conscious VE was not driven by individuals with higher task performance. Dashed lines represent 95% confidence intervals on the regression line. The right panel shows the VE for subjectively invisible and visible trials in a subset (n = 21) of participants whose flash localization accuracy on invisible trials indicated substantial support (BF_null_ > 3.33) for the null hypothesis of chance performance. Together these analyses suggest that the non-conscious ventriloquist effect holds even for objectively chance levels of performance. Error bar are 95% confidence intervals.

As an additional control, we sought to screen subjects who may have been above or below chance levels of performance when they rated the flash as invisible. To do this we ran a Bayesian binomial test on each participant’s accuracy on invisible trials to see if there was evidence for or against the null hypothesis of the data coming from a binomial distribution with success probability of 0.5 (i.e., 50% accuracy). Only participants with a BF substantially favoring the null hypothesis of chance performance (BF_null_ > 3.33) were retained and the main VE analysis was repeated on this subset. We found 21 of our 30 participants had evidence in favor of the null hypothesis of chance performance. Importantly, using this subset of participants still produced a clear but slightly reduced invisible VE of 1.52 deg. (0.19) that remained statistically significant (*t*(20) = 7.872, *p* < 0.001) and provided decisive evidence for a non-conscious VE (BF_alt_ > 999).

### EEG Signals of Flash Location Processing

We decoded the left/right location of the flash from 64-channel EEG signals using multivariate LDA in a subset of 20 subjects who were at chance at discriminating the flash location on invisible trials (Figure 5A). We found that invisible flash location was significantly decodable from 84 to 224 ms (cluster *p* = 0.0002), indicating preserved early, putatively feed-forward spatially-selective processing during unconscious masked trials. On visible trials, however, decoding was sustained longer with one early cluster spanning from 112 to 216 ms (cluster *p* = 0.0018) and another later cluster from 248 to 384 ms (cluster *p* = 0.0004). Lastly, visible trials were associated with significantly higher decoding accuracy from 308 to 328 ms (cluster *p* = 0.0074), indicating that the neural representation of visual spatial location varied with awareness only after relatively late (>300 ms) processing.

**Figure 5.**
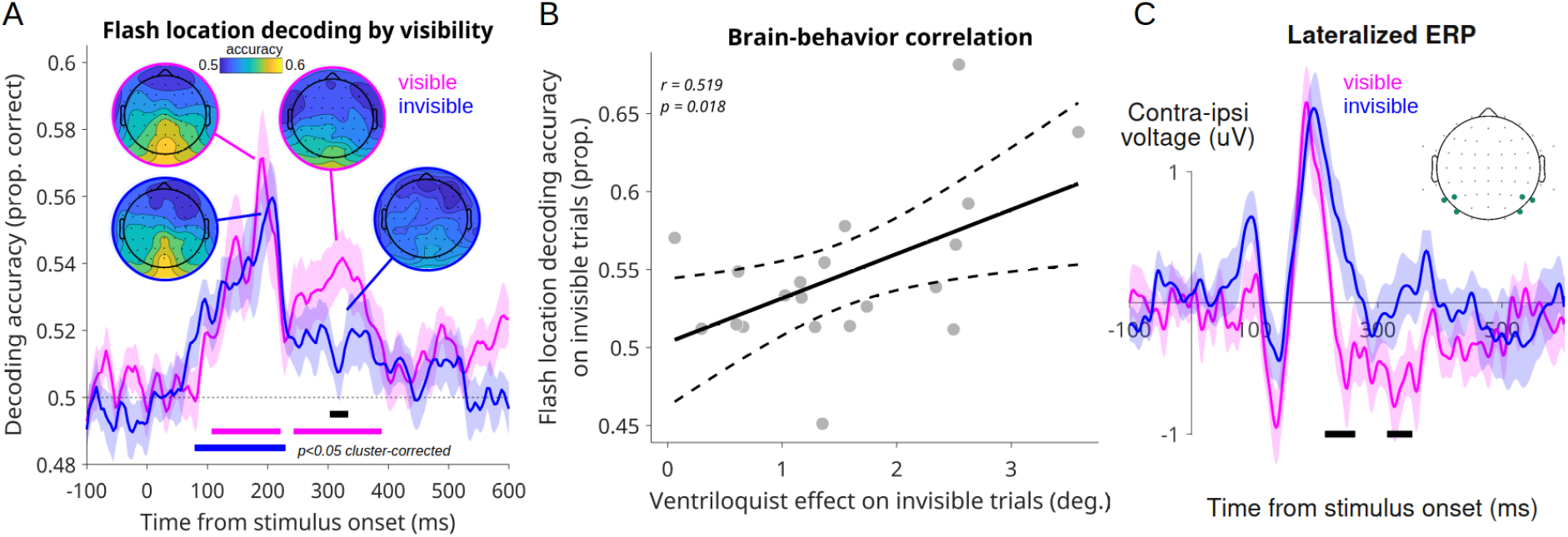
EEG representations of visual stimulus location in chance subjects and relation to behavior (n = 20). **A)** Shows EEG decoding accuracy over time for a classifier trained to predict the location (left/right) of the flash stimulus using all EEG channels, separately for visible and invisible trials. Magenta squares below show significantly above-chance timepoints (cluster-corrected) for visible trials, blue squares show invisible trials, and black squares show the difference between visible and invisible. Initial decoding accuracy was comparable for visible and invisible trials but only visible trial decoding persisted beyond ~220 ms and significantly differed from invisible trials around 300 ms. Results are consistent with a preserved early, feed-forward sweep of processing during invisible trials but a lack of sustained “broadcast” of non-conscious information. Shaded regions denote the SEM. Inset topographical maps show decoding accuracy across the scalp from a searchlight procedure using visible and invisible trials and early (170-200 ms) and late (230-360) EEG signals. **B)** The cross-subjects correlation between early decoding accuracy (170-200 ms) on invisible trials and the VE on invisible trials. **C)** Lateralized ERPs from electrodes contralateral minus ipsilateral to the visual stimulus on visible and invisible trials. In line with the decoding results, early lateralization of brain responses caused by the location of the visual stimulus were comparable across visible and invisible trials and only differed after 220 ms (black squares denote cluster-corrected differences between visible and invisible ERPs). Shaded regions denote SEM and inset topography shows lateralized electrodes used to generate ERP waveforms.

To better understand the topographical features contributing to flash location decoding, we performed a searchlight decoding analysis across the entire EEG channel space. Decoding accuracy at each searchlight location for the early decoding peak (170-200 ms) showed strongest accuracy at midline and bilateral occipital and occipital-parietal electrodes (see inset Figure 5A). The early decoding topographies were highly similar for visible and invisible trials, suggesting similar neural mechanisms underlying early conscious and non-conscious location processing. During later processing however (230-360 ms), the topographies diverged, with no electrodes showing clear decoding on invisible trials but a similar set of posterior electrodes showing sustained decoding on visible trials. Although we cannot infer the neural generators underlying these decoding effects, they seem to suggest that conscious location information is represented in patterns of sustained posterior scalp EEG activity.

Together these findings indicate that location information on invisible trials was processed comparably to visible trials during early stimulus processing, but were not processed to the point of global “broadcast”, thought to occur around 250 – 300 ms (Del Cul et al., 2007; Salti et al., 2015; Sergent et al., 2021). This suggests that any multisensory integration occurring on invisible trials may take place before 250 ms (i.e., before conscious broadcast according to GNWT). If this is true, then one might expect that the strength of the early decoding signal on invisible trials ought to predict the strength of the VE on those same invisible trials. We tested this by correlating decoding accuracy averaged across the early time window (170–200 ms) on invisible trials with the size of the invisible VE across participants (Figure 5B). This revealed a significant positive correlation (*r* = 0.519, *p* = 0.018), indicating that subjects with stronger earlier decoding on invisible trials had a larger unconscious VE. This finding lends correlational support to the notion that early location representations (~200 ms), despite being invisible, may be sufficient for audio-visual spatial integration.

Finally, we examined more traditional event-related potential (ERP) markers of visual processing to see whether the decoding results generalized to simpler analyses. To this end, we isolated ERP responses specific to the location of the visual stimulus by computing ERPs contralateral and ipsilateral to the visual stimulus and then computing subtraction waves to examine spatially-specific neural activity as a function of visibility (Figure 5C). Note that this subtraction also inherently removes mask-related neural activity since the mask was equally present regardless of stimulus location. We found that early lateralized ERP responses did not significantly differ between visible and invisible trials, in alignment with the early decoding results. However, two significant later clusters emerged such that a larger lateralized negativity from 220 to 258 ms (cluster *p* = 0.011) and again from 320 to 350 ms (cluster *p* = 0.026). Thus, ERP analysis of lateralized visual activity suggest that subjective visibility is associated with relatively late spatially-specific processing in this task.

## Discussion

We investigated the role of conscious awareness in a well-studied multisensory integration phenomenon known as the Ventriloquist effect (VE). We found that visual stimuli rendered non-conscious via metacontrast masking nevertheless biased auditory perception towards them. Visual stimuli could not be localized better than chance levels yet our results show that they were still integrated into auditory perception, revealing a clear non-conscious VE. Moreover, we found that EEG decoding of invisible flash location showed an initial representation comparable to that of seen stimuli, but was not sustained beyond ~220 ms, indicating that the non-conscious integration may have occurred prior to 220 ms, before global broadcast is thought to occur according GNWT. In line with this, early decodability predicted the size of the non-conscious VE across participants.

Our results add to previous evidence suggesting that integration of non-conscious inputs is more pronounced than once believed, particularly in the domain of multisensory integration. Indeed, recent findings indicate that associative learning can occur between two non-conscious stimuli in different sensory modalities, as measured by RT priming effects (Scott et al., 2018). Another study found semantic priming effects on RT for masked audio-visual primes (Faivre et al., 2014). Our results critically extend these findings, however, by going beyond RT effects to show that an unconscious stimulus in one modality actually alters the conscious perception of a stimulus in another modality. In other words, our findings suggest that non-conscious multisensory integration actually shapes the contents of conscious experience and not only produces priming effects in RT tasks.

In this vein, our findings are concordant with two recent studies showing similar effects. First, the (Delong et al., 2018) study that directly motivated ours found a similar VE to invisible flashes using dCFS as a means of suppressing flash awareness. Additionally, a recent study found that motion after effects caused by adaptation to a continuous motion signal could be boosted by a congruent auditory motion stimulus and that this effect was observable even when the adapting visual motion stimulus was rendered non-conscious using CFS (Park et al., 2024). Thus, while both studies come to the same general conclusions as ours, it should be noted that CFS has received criticism as a tool for studying unconscious processing due to the possibility (and some evidence) that observers may briefly be aware of low-level features of the suppressed stimuli but simply confuse them as part of the mask, which typically varies across trials along many dimensions of color, shape, size, luminance, etc. (Gelbard-Sagiv et al., 2016) Thus, our results add important evidence in favor of non-conscious multisensory integration using a different masking approach less susceptible to those concerns.

Additionally, the decoding analysis of concurrent EEG signals helps to pinpoint the general time frame of non-conscious AV spatial integration in the VE. If we accept that the unconscious AV integration likely occurs while there is still an active representation of the visual stimulus, then our our data indicate that this may occur within the first 220 ms of processing, which is when we ceased to be able to decode the unconscious flash location. This claim is further supported by our finding that flash location decoding performance during this early time window correlated with the size of the unconscious VE in the behavior. During normal conscious multisensory perception, however, prior studies suggest that integration-related neural dynamics may persist for up to ~700 ms (Rohe et al., 2019), suggesting that consciousness may continue to play a role after initial unconscious integration. Although we could not measure a neural signature of integration directly due to poor decoding performance of the sound location (see Methods), this would be an important step for future research.

It is also worth noting that the magnitude of the non-conscious VE we observed was rather large, comparable even to that of the visible flash trials. This departs from the finding of Delong et al. (2018) who observed an invisible VE of ~20% of the visible one, compared to about 82% in the current study. One possible explanation for the large effect observed here is that metacontrast masking may preserve more unconscious processing compared to CFS, which may suppress information processing at an earlier stage. Indeed, previous work has suggested an advantage for meta-contrast masking over CFS in affording unconscious perceptual processing (Peremen & Lamy, 2014). Another possibility is that the conscious VE in our data was somewhat minimized since even when participants did report seeing the stimulus, the most frequency PAS rating was a one or “brief glimpse” (see Figure 2B). This suggests that perhaps greater awareness of the stimulus (e.g., PAS = 2 or 3) may be associated with an even larger VE, however we had too few trials in those categories to meaningfully test this.

Our study also has several limitations. First, we did not measure objective flash localization during the main AV task, only auditory localization and subjective visual awareness. This choice was made to minimize the cognitive demands of answering three question on each trial and also avoid erroneous responses due to confusion or working memory limitations. Although we do show that trials rated as invisible were not localized better than chance both before and after the main AV task, not having objective measures of flash discrimination during the AV task leaves open the possibility that stimuli were not invisible by objective standards during this task.

Second, we were unable to decode the sound location, which limited the conclusions we could draw from the neural data regarding AV integration. Other studies have successfully decoded sound location from EEG (Bednar et al., 2017), but typically for sounds spaced much further apart (e.g., 90 deg. apart) than the 5 or 10 deg. separation used here. Third, by aiming to render visual stimuli invisible, our study ultimately compares unconscious perception to very weak conscious perception (see PAS rating distributions in Figure 2B, for instance). This is also evident in our decoding results, which show only minor differences in the location information present between visible and invisible trials (see figure 5). Thus we cannot claim that consciousness has not role in AV integration, just that it is seemingly not required. The same can conclusion can be made about the role of top-down spatial attention, which is presumably absent on invisible trials. While this shows that top-down attention is not necessary for the VE, it does not preclude a role for top-down attention in modulating the VE. Finally, our task differed from most typical VE paradigms by having participants report the visual stimulus as well as the auditory one, thereby focusing more attention on the visual stimulus. Although attention likely has little impact on invisible trials, it would be interesting to see if comparable unconscious effects could be observed without awareness and attention.

Although our study was inspired by predictions made by the GNWT of consciousness, our results provide constraints on any approach to consciousness that emphasizes integrative functions of consciousness, such as the Integrated Information Theory (IIT; see Noel et al., 2019 for a conncetion between IIT and multisensory integration). With respect to GNWT, however, we can see several ways the theory might adapt in light of these findings. It could be that some integration is possible unconsciously but not all forms of integration. Specifically, it could be the case that relatively low-level (i.e., spatial or temporal congruency-based) forms of integration, such as in the VE, can occur either subcortically or via direct auditory-visual cortico-cortico projections (Ghazanfar & Schroeder, 2006). However, perhaps some forms of integration, such as those based on semantic congruency could require higher-level processing only possible if stimuli are consciously perceived and activate global workspace neurons. This is indeed suggested by recent findings showing that AV integration based on semantic cues does not seem to occur when stimuli are presented non-consciously (Delong & Noppeney, 2021). Similar arguments have been made on the basis of the McGurk effect seeming to depend on conscious perception of the visual component of the stimulus (Palmer & Ramsey, 2012). Thus, while it might be the case that certain forms of multisensory integration require conscious perception, it seems clear that not all forms do. Our findings begin to constrain the nature and timing of multisensory interactions in the absence of awareness, putting pressure on integration-based theories of consciousness to better specify the role of consciousness in multisensory processing.

## Notes

**Conflicts of interest:** none

### Competing Interest Statement

The authors have declared no competing interest.

### Summary of Updates

Expanded EEG and behavioral analysis, additional descriptive statistics, and updated discussion.

